# Improved inference of chromosome conformation from images of labeled loci

**DOI:** 10.1101/409847

**Authors:** Brian C. Ross, James Costello

## Abstract

We previously published a method that infers chromosome conformation from images of fluorescently-tagged genomic loci, for the case when there are many loci labeled with each distinguishable color. Here we build on our previous work and improve the reconstruction algorithm to address previous limitations. We show that these improvements 1) increase the reconstruction accuracy and 2) allow the method to be used on large-scale problems involving several hundred labeled loci. Simulations indicate that full-chromosome reconstructions at 1/2 Mb resolution are possible using existing labeling and imaging technologies. The updated reconstruction code and the script files used for this paper are available at: https://github.com/heltilda/align3d.

---

Measurement of *in vivo* chromosome conformation is a major unsolved problem in structural biology despite its known biological importance [1]. The present state-of-art is either to obtain indirect information about conformations using 3C-derived methods which measure DNA-DNA contacts (typically in a cell-averaged population) [2] or else to directly measure the cellular locations of individual chromosomal loci in single cells by microscopy [3] The major limitation of direct localization is one of throughput: only ∼3 — 5 labeled loci can be uniquely identified ‘by color’ in a standard microscope image, whereas a whole-chromosome reconstruction would involve labeling and identifying hundreds or thousands of loci. Several research efforts aim to remove this limitation using either experimental or computational approaches. The experimental approaches aim to allow an increased number of labels that can be distinguished in an image [4-6]. Alternatively, computational methods are being developed to infer the identity of each label if it cannot be uniquely identified in an image [7-9], by using the known label positions along the DNA contour. Here we focus on the computational inference method called `align3d` Ref. [8], and present improvements to allow high-quality, chromosome-scale conformational reconstructions.

First, we briefly describe the `align3d` algorithm. Using a) the genomic locations and colors of labeled loci and b) the spatial locations and colors of spots in a microscope image, together with a relation tying the genomic distance between two loci to their average spatial displacement, this method constructs a table of ‘mapping probabilities’ *p*(*L* → *s*) for a given labeled genomic locus *L* having produced spot *s* in the microscope image. Each mapping probability *p*(*L* → *s*) is calculated by dividing the summed statistical weights of conformations where locus *L* maps to spot *s,* which we term a mapping partition function and denote *Z*_*L* →_*_s_*, by the full partition function *Z* that is the summed weight of all conformations. A proper calculation of *Z*_*L* →_*_S_* and *Z* would consider all conformations having no more than one locus at any given spot in the image^1^, similar to a traveling salesman tour [10], but this exact calculation is intractable for large problems. Instead, `align3d` counts all conformations for which *adjacent* loci do not overlap at the same spot (see Figure 1), using a variant of the forward-backward algorithm [11] that can propagate between non-adjacent layers. This is a major source of error as the vast majority of conformations contributing to the partition function overlap at non-adjacent loci, and one consequence is that the normalization of mapping probabilities makes no sense for a non-overlapping conformation, as Σ*_L_P*(*L* → *s*) can exceed 100% for certain spots. To recover from this error, `align3d` assigns a penalty to each spot and iteratively adjusts these penalties until the spot normalization is sensible. Although somewhat ad hoc, use of spot penalties recovers significant information about medium-sized conformations (∼ 30 labeled loci), although larger simulated experiments (∼ 300 loci) have convergence problems due to the cost function plateauing at very small or large values of the spot penalties.

**Figure 1:**
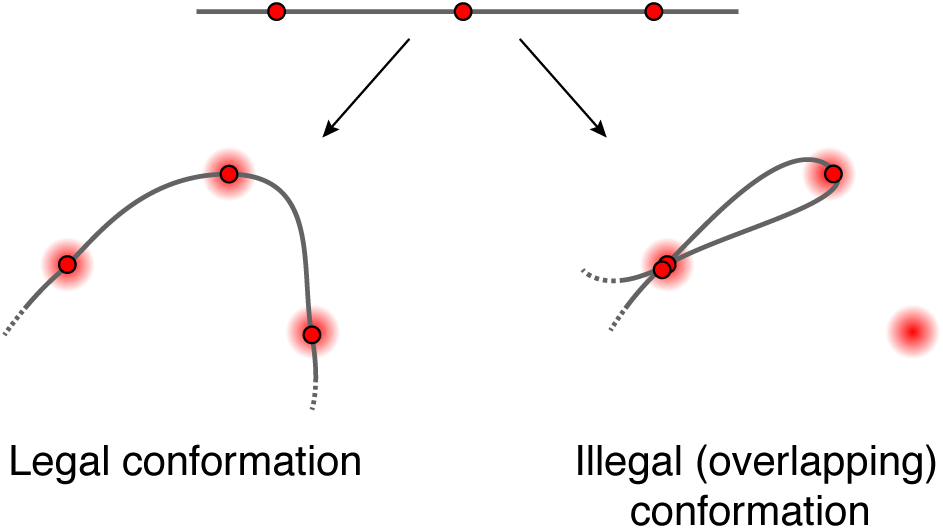
Canceling out overlapping conformations. Schematic showing one legal and one illegal conformation passing through spots *A, B* and *C.* `align3d` counts both legal and overlapping conformations in estimating the partition *Z* (although it is able to prevent *adjacent* loci from overlapping).

The final step is to use the mapping probabilities to construct the range of likely conformations compatible with the microscope image. Uncertainty in the conformation results from inaccuracy or uncertainty in the mapping probabilities due to three factors: inaccuracy in the DNA model (the relation between genomic and spatial distance), error in estimating the partition functions, and the inherent uncertainty in the data even with a perfect reconstruction algorithm. Improvements to the DNA model will require direct measurements of *in vivo* conformations, which have not yet been made. Here we focus on improving the partition function estimate, using two different strategies. First, we give an efficient method for optimizing the spot penalties when there are hundreds of spots in the image. Next, we provide formulas for the partition functions which allow them to be estimated to arbitrarily high accuracy (given enough computational time), without using spot penalties or any optimization. As we show using simulations, these two methods used individually or in tandem permit confident, chromosome-scale conformational reconstructions using existing experiments.

## Methods

First we provide a method for efficiently optimizing the spot penalties regardless of the number of labeled loci. This rule guarantees that a) the rate of missing spots is as expected, and b) the mapping probabilities are properly normalized. Let *q*_*s*_ denote the penalty attached to spot *s*; then the update rule for that spot penalty is:

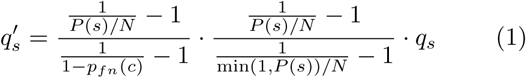

where *N* is the number of loci, *P(s*) = Σ*_L_ p*(*L* → *s*) is the total probability of mapping any locus to spot *s,* and *Pf*_*n*_(*c*) is the estimated rate of missing spots having color c. The justification for this rule is given in Appendix 1.

We can also update a penalty 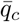 that is associated with *missing* spots of color *c*. This gives a faster way to enforce a desired missing spot rate because there are fewer 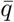 penalties than 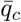 penalties. An update to is equivalent to a reverse update to all *q*_*s*_ for spots *s* of color *c*, so the update rule is:

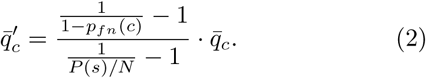

Typically, we first optimize the 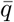 parameters to achieve a target missing spot rate, then optimize the *q* parameters to enforce *P*(*s*) ≤ 1 while maintaining the missing spot rate. In either case, we apply Eq. 1 or 2 to bring the *q* or 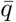 parameters close to their final values. When the cost function stops improving, we switch to the steepest-descent algorithms used in Ref. [8] to polish *q* or 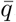.

Next, we give two exact formulas for the partition functions *Z* _*L* ⟶ *s*_ and the full partition function *Z* that determine our locus-to-spot mapping probabilities. We focus on the full partition function *Z* since the formulas for *Z*_*L*⟶ *S*_ are identical. The largest term in each formula, which we denote 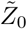 (or 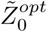 when spot penalty optimization is used), is the original estimate from Ref. [8] calculated using a variant of the forward-backward algorithm [11]. Additional terms are computed in the same way, except that certain loci are constrained to map to certain spots. All of the constraints we will apply are *illegal constraints*, in that they force multiple loci to overlap at some spot in the image; therefore these terms only count illegal conformations that we would like to remove from the baseline calculation. By computing these terms and subtracting them from 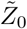 we eliminate the overlapping conformations and improve the calculation. It turns out that this process erroneously subtracts conformations with multiple overlaps more than once and thus we have to add back in higher-order corrections (i.e. partition functions having multiple constrained spots). Repeating this logic leads to exact formulas for *Z* taking the form of series expansions, which are dominated by the lowest-order terms as those have the fewest restrictions on conformational overlaps. Figure 2A illustrates an example of such a series expansion, where each parenthetical subscript (*XY…*)_*s*_ on a term label denotes an illegal constraint forcing loci *X, Y*,… to overlap at spot *s* when that term is calculated. We use this notation throughout.

**Figure 2:**
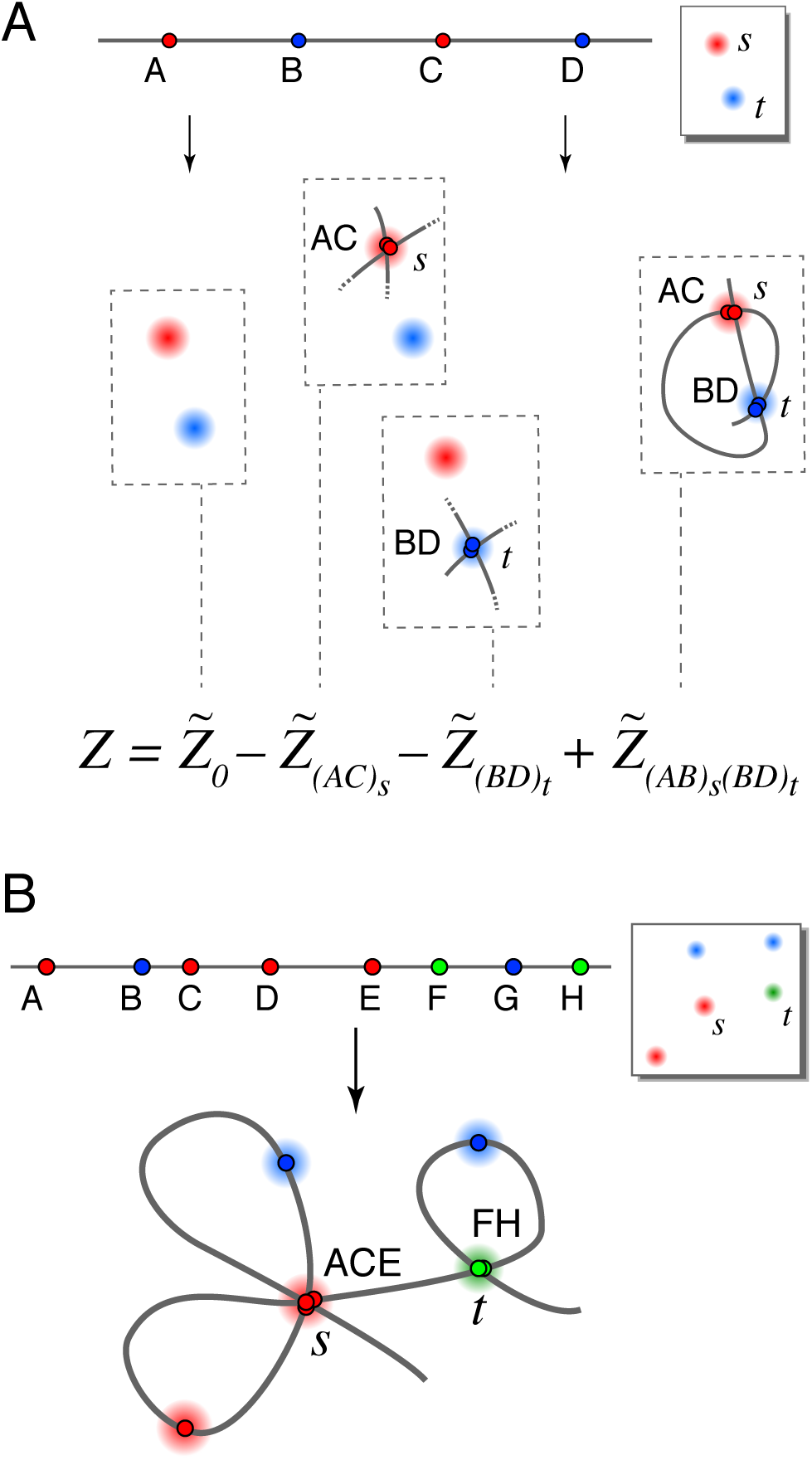
Series expansions. **A.** Schematic showing terms in a series expansion, in a case where series 1 and series 2 have the same terms. The full series gives the exact partition function for the 4-locus experiment shown where only 2 spots appeared in the image (due to a high rate of missing spots). Cartoons show only the constrained loci for each term (so for example each term includes the illegal conformation visiting spots *s* →t → *s* →*t*). **B.** An illegal conformation for which loci *A, C* and *E* overlap at spots *s,* and loci *F* and *H* overlap at spot *t.* Series expansion 1 includes this conformation in terms 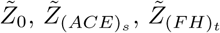, and 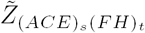. Series expansion 2 includes this conformation in the same terms with the addition of 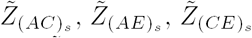, and 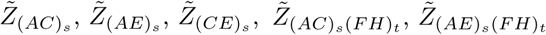.

There are two ways we might remove conformations containing overlapping loci, leading us to two different series expansions for the true partition function *Z.* Suppose that we are calculating the term 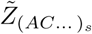 whose single illegal constraint forces loci *A,C,…* to overlap at spot s. One option is to forbid any of the other unconstrained loci from also mapping to spot *s,* since spot *s* is already overused. This leads to series expansion 1. Alternatively, allowing further overlaps with spot *s* from the unconstrained loci gives us series expansion 2. Figure 2B illustrates the differences between the two series.

Each of the two series expansions is a weighted sum over *all possible illegally-constrained terms* having two properties: 1) each locus and each spot appear at most once in the indices, and 2) two or more loci map to each constrained spot. To be formal, we use Ω to represent the set of all possible illegal constraints: each element of Ω, consists of a set of two or more non-adjacent loci and a single spot where they are forced to overlap. Each expansion thus takes the form

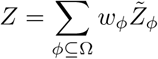

where *Z* _*ϕ*_ is zero if any two constraints share a locus or spot. We will choose the integer weights *w* _*ϕ*_ so as to cancel out the overlapping conformations. By symmetry arguments, the weighting factor should not depend on the identities of the loci or spots, but rather only by the number of constrained spots *n*_*ϕ*_, and the number of loci 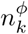 involved in each *k*^*th*^ constraint. For example, *w(ACE)*_*s*_*(BD)*_*t*_ is determined by *n*_*ϕ*_ = 2, 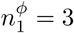 and 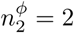.

Here we specify each series expansion by giving a formula for the weights *w*_*ϕ*_ in terms of *n*_*ϕ*_ and the various 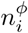 We also explain how to select an appropriate set of terms *ψ* when there are too many terms to evaluate. Our selection prohibits any legal or overlapping conformation from contributing a negative weight to the partition function estimate, thereby guaranteeing positive mapping probabilities and allowing use of the reconstruction-quality metrics given in Ref. [8]. Derivations of the coefficient formulas and the term-selection criteria for each series expansion appear in Appendix 2.

### Series expansion 1

For series expansion 1, we do not allow the unconstrained loci to map to spots that were used in constraints. Then the weights *w*_*ϕ*_ are given by:

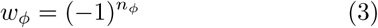

To select terms for a series approximation, we first choose a set of illegal constraints *ψ* to disallow, then include all series terms *Z*_*ϕ*_ containing only those constraints: i.e. *ϕ* ⊆ *ψ*. This guarantees non-negative mapping probabilities. In order to efficiently evaluate the largest terms, we recommend selecting the *N*_*ψ*_ constraints having the highest product of mapping probabilities in the baseline calculation 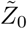 (or 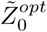 if spot penalties will be used). For example, we would include (*AC*)_*s*_ if *p*(*A* →*s*) • *p*(*C* →*s*) is sufficiently large.

### Series expansion 2

For series expansion 2, the unconstrained loci are allowed to map to spots that were used in constraints. Then the weights *w*_*ϕ*_ are given by:

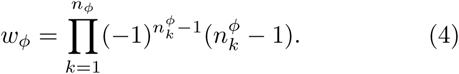

To select terms for a series approximation, we first choose a set of *N*_*ψ*_ singie-locus-to-spot mappings Ψ, then include all terms *Z*_*ϕ*_ whose illegal constraints use only mappings in For example, the constraint (*AC*)_*s*_ would be included if Ψ ⊆ {*A* ⟶ *s*, *C* ⟶ *s*}. In order to select the largest terms, we recommend building Ψ from the *N*_*ψ*_ largest mapping probabilities calculated from 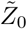 or 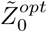.

## Results

We tested the improved `align3d` method by generating random chromosome conformations and simulating the process of error-prone labeling, imaging and finally producing the locus-to-spot mapping probabilities. We considered three scenarios for our simulations. 1) The ‘Toy’ scenario involves 10 genomic loci, where each locus is labeled using one of 3 colors. For these simple problems the partition function can be calculated exactly. 2) Our simulated Experiment 1 uses standard DNA labeling methods and traditional 3-color microscopy to label 30 loci with 3 colors, thus interrogating a significant fraction of a chromosome contour. 3) Our simulated Experiment 2 labels 300 loci across a chromosome-length contour. The reconstruction of Experiment 2 is made possible by using the Oligopaints labeling technique [4] to label in 20 different colors.

For each scenario, we randomly generated 100 conformations using a wormlike chain model (packing density = 0.3 kb/nm, persistence length = 300 kb, as suggested by the measurements of Ref. [12]); applied a random labeling at a mean density of 1 locus per megabase; and simulated experimental error: 100/200-nm Gaussian localization error in xy/z, a 10% rate of missing labels, and a 10% rate of nonspecifically-bound labels. A typical simulated experiment from the Toy scenario is shown in Figure 3A.

**Figure 3:**
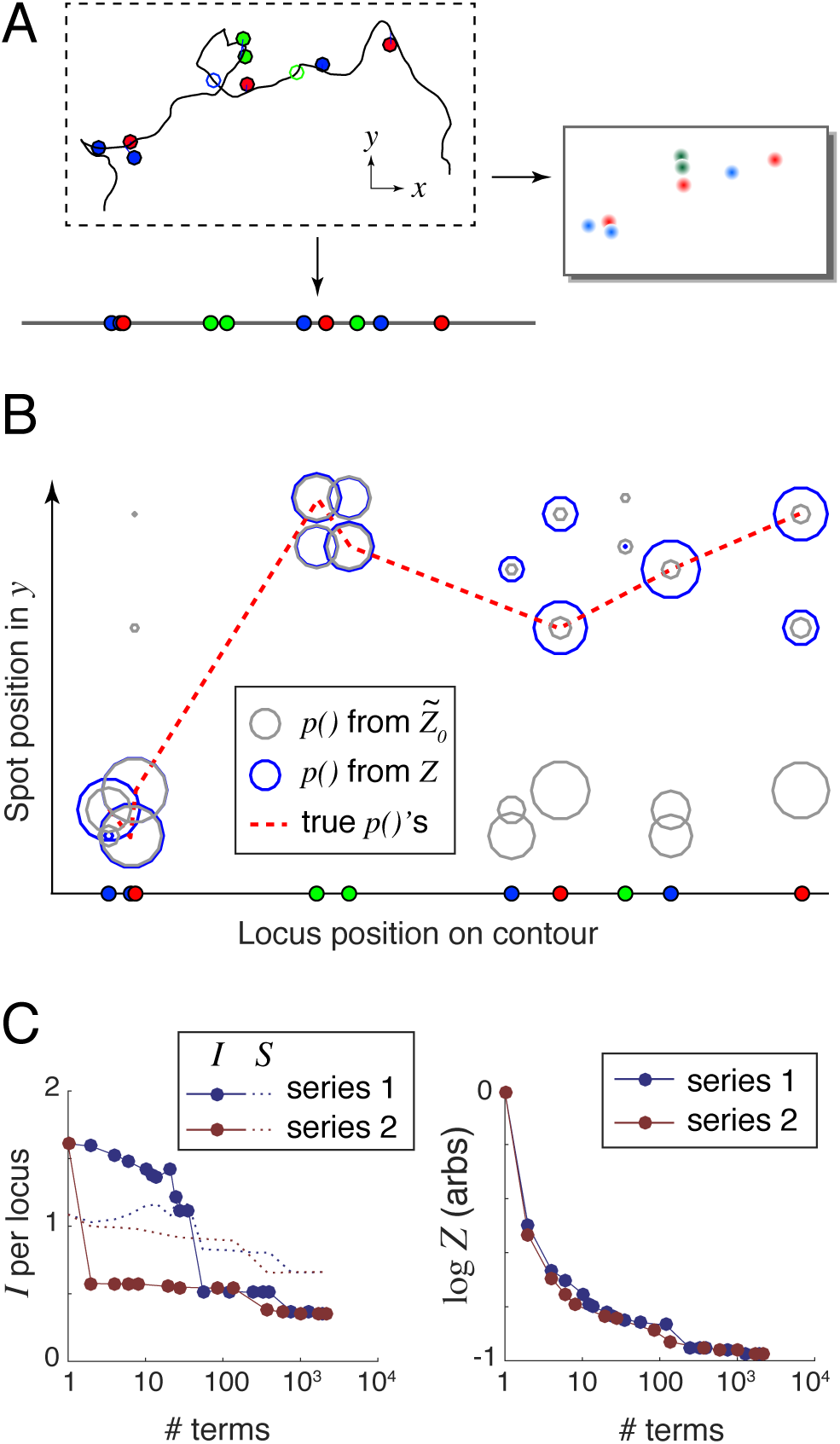
Example reconstruction. **A.** Randomly generated and labeled chromosome contour with simulated experimental error: localization error (lines offsetting spots from the labeled genomic loci) and missing labels (open circles). This example lacks nonspecifically-bound labels (floating spots). **B.** Spot mapping probabilities calculated using both the largest series term 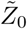 (grey circles), and the exact *Z* that can be computed using 2210 series terms (blue circles). The dotted red line connects the true locus-to-spot mappings, which are used to calculate the unrecovered information. *I*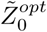 bits/locus and *I(Z)* = 0.54 bits/locus. C. Unrecovered information *I* and entropy *S* (left panel) and log *Z* (right panel) versus the number of terms used in the series expansions.

For each simulated conformation, we fed the label positions and colors together with the simulated 3D images into the `align3d` algorithm to produce locus-to-spot mapping probabilities. For example, the experiment shown in Figure 3A produced the mapping probabilities shown graphically in Figure 3B using circles, where the size of each circle indicates probability magnitude. Here grey circles show the mapping probabilities computed from 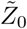 with no use of spot penalties, and blue circles show those same probabilities computed using the exact *Z.* This example shows how excluding high-weight and heavily-overlapping conformations reduces and improves the partition function estimate (see Figure 3C) and concentrates the probability mass into the ‘true’ locus-to-spot mappings (shown connected by the dotted red line in Figure 3B).

Our reconstruction quality metric is the amount of *unrecovered information* from the mapping probabilities, defined as *I* = – ⟨log *p*(*L*_*i*_ ⟶ *S*_*i*_) *⟩*_*i*_ where the average ⟨·⟩ is taken over the set of true locus-to-spot mappings (*L_i_, s_i_*). The ideal case of *I* ⟶ 0 implies a perfect reconstruction with no mistakes and zero uncertainty, but in practice *I* is always positive. In a real experiment where the true mappings are not known, we use a proxy for unrecovered information that we term entropy, defined as *S* = – ⟨*p(L*_*i*_ *⟶ S*_*j*_) log *p(L*_*i*_ *⟶S*_*j*_*) ⟩*_*ij*_ whose average is taken over all locus-to-spot mappings, not just the correct mappings. The goal is to have *S ≈ I* so that a real experiment will have an accurate estimate of the reconstruction performance. The left-hand panel of Figure 3C shows how *I* and *S* depend on the accuracy of the calculation for the simple example shown, using either of the two series expansions and varying the number of terms from 1 (simply 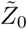) to 2210 which is the full set of terms for either series and thus computes *Z* exactly. Entropy generally overestimates the amount of unrecovered information (see Supplementary Figures S1 and S2), because the large mapping probabilities should be even larger, and the small ones even smaller, than their assigned values (see Supplementary Figure S3). We believe this miscalibration is mostly caused by `align3d` overestimating the missing spot rate.

### Validation of Equations 1-4

We first validated each of the two series expansions by comparing them against exact partition function calculations for the simulated Toy experiments. In all cases, both series expansions, when taken to their maximum number of terms, exactly reproduced the partition function calculations obtained by direct enumeration over all possible non-overlapping conformations. This test validates Equations 3 and 4. We also verified that both series expansions could be used in conjunction with spot penalty optimization (Equations 1 and 2), both by numerically validating the cost function gradient calculation and by testing for convergence on these small problems.

### Improved optimization allows large-scale reconstructions

First, we tested whether the iterative spot-penalty optimization rules given by Eqs. 1 and 2 could work on large-scale problems such as those of Experiment 2, where the old gradient descent optimizer in `align3d` had difficulty [8]. The results are shown in Figure 4, which compares the number of iterative steps required to converge the 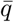 (missing-spot penalty) and *q* (spot penalty) parameters without/with use of our improved optimization rules (labeled ‘old’/‘new’ respectively in the legend). Since the spot penalties *q* are optimized for probability normalization only after 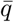 parameters have been optimized to achieve a desired missing spot frequency, we only attempted to optimize the *q* parameters for simulations where 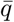 converged. There were two results from this experiment. First, more attempts to optimize the 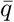 and *q* parameters successfully converged when using the new optimization rules in conjunction with gradient descent, as indicated by the greater volume of the ‘new’ histogram and the correspondingly larger numbers shown in the legends. Secondly, of the trials that did converge, our new method required significantly fewer iterations and thus less computation time than the old method, as indicated by the relative skews of the distributions.

**Figure 4:**
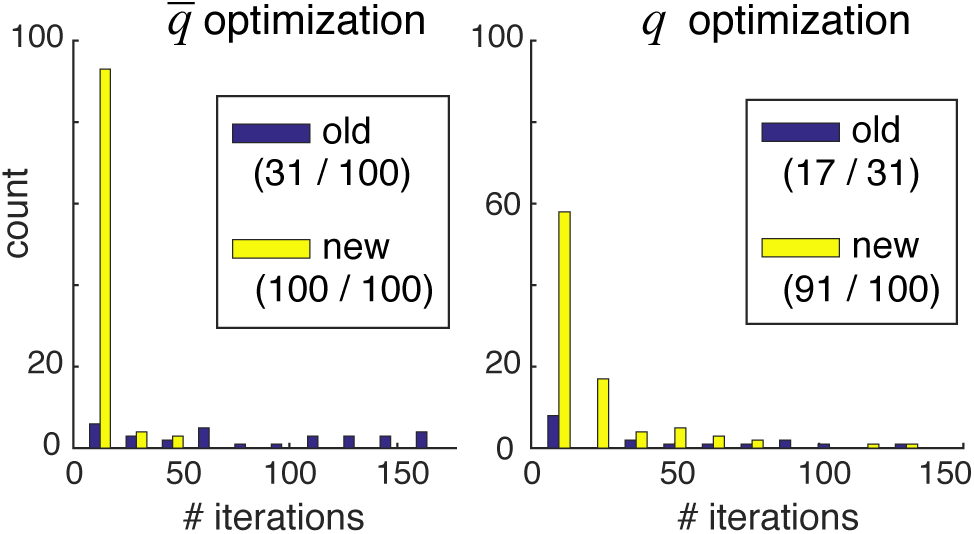
Comparison of old and new optimization methods. Each panel compares the number of iterations required to achieve convergence using the old (purple) versus new (yellow) optimization methods. Only trials that successfully converged are counted, so the histograms are not normalized relative to each other. The first number in parentheses of each legend entry shows the number of converged trials, and the second number shows the total number of trials. Note that the second numbers in the right-hand panel equal the first numbers in the left-hand panel, since we required convergence in 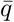 in order to attempt optimization of the *q* parameters.

### Use of more colors dramatically improves reconstructions

Our most striking result is that simulations of the ambitious Experiment 2 produce far better results than even the Toy scenario, despite the fact that these simulations have more loci per color than either the Toy scenario or Experiment 1. This can be seen in the amount of unrecovered information *I* shown in the simulation-averaged plots of Fig. 5A. Thus a push to 20-color labeling could prove critical for genomic reconstruction at the chromosome scale and beyond. At the end of this section we revisit Experiment 2, in order to assess the reconstruction quality when analyzing more realistic DNA contours having tighter confinement.

**Figure 5:**
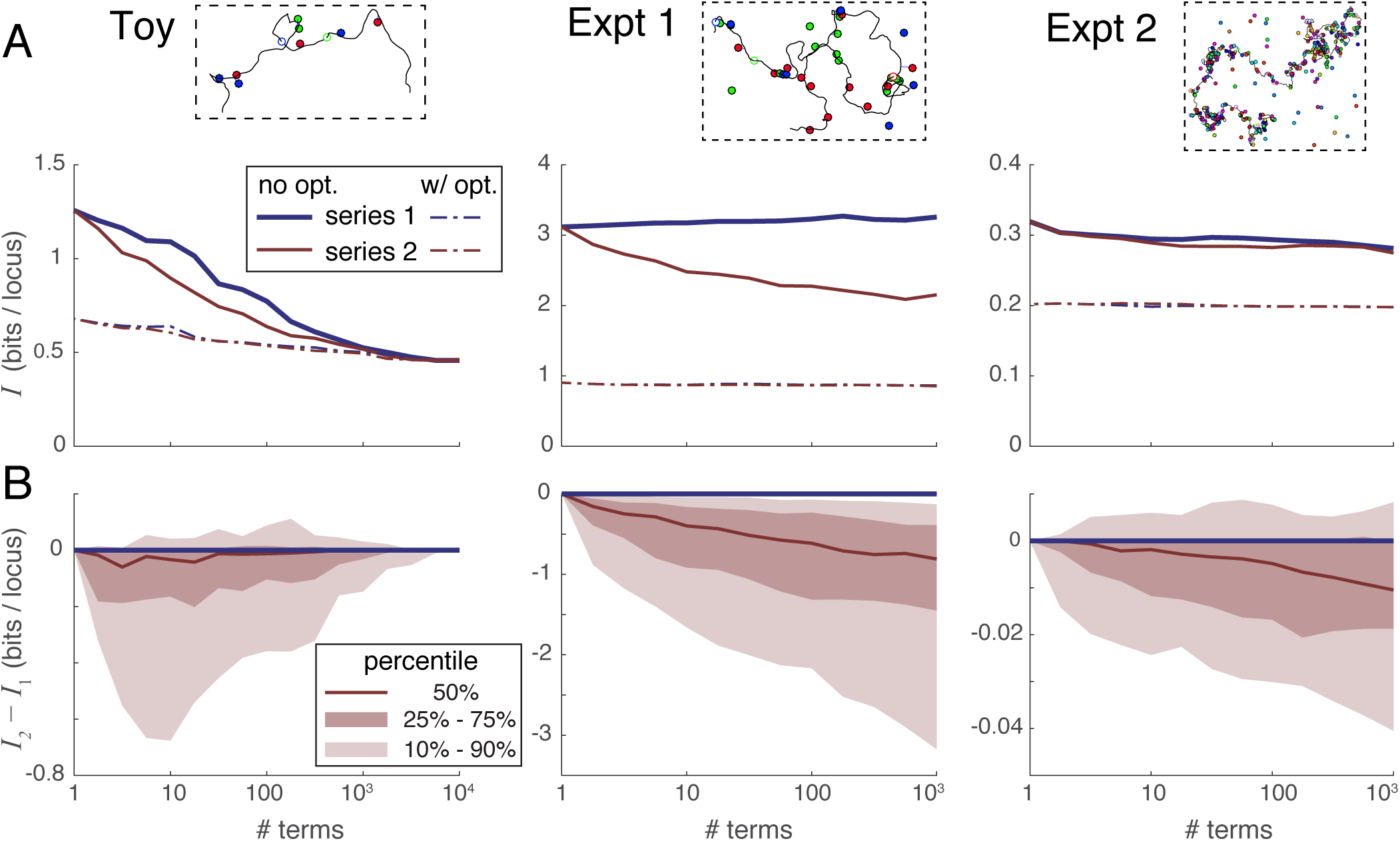
Comparison of the convergence rates of series expansion 1 and series expansion 2. **A.** Median unrecovered information *I* as a function of the number of terms used in each series expansion, without using spot penalty optimization (solid lines) versus with optimization (dotted lines), and over the three simulation scenarios (panels left-to-right). Each curve was derived from the 100 individual curves corresponding to the 100 simulations in each scenario using a simple point-by-point median average. **B.** Percentile distribution of the difference between the unrecovered information using series 2 minus the unrecovered information using series 1; the fact that this difference quickly drops below zero in nearly all individual simulations shows that series 2 recovers more information in the first few terms than does series 1.

### Series expansion 2 outperforms series expansion 1

Next, we compared the convergence properties of our two expansions on the three scenarios of simulated experiments. Figure 5A gives a sense of how the amount of unrecovered information varies with the number of terms taken in each series, without (solid lines) and with (dotted lines) the use of spot penalties. Each of the 3 panels summarizes all 100 simulated experiments of that scenario, and each experiment in that scenario shows a unique relationship between information recovery and number of series terms computed. Representative curves of individual experiments in each scenario are shown in Supplementary Figure S1. In order to summarize these very dissimilar curves, Figure 5A shows a median average of all 100 individual experimental curves taken at each data point. Note that this averaging process does not necessarily preserve the shape of the curves from typical individual simulations.

In order to directly compare the two series expansions, we plotted their difference in unrecovered information *I*_*2*_ *— I*_*1*_ versus the number of series terms in Figure 5B. In this case, we plotted the full distribution showing the median (50th percentile) as well as the 10th, 25th, 75th and 90th percentile curves. These plots show directly that series 2 almost always outperforms series 1 when only a few terms can be evaluated. The reason is that the terms in series 2 are larger in magnitude owing to their looser constraints, and thus remove the extraneous part of the partition function more quickly than the terms of series 1 (see Supplementary Figures S1 and S4). Based on these results, we recommend using series expansion 2 in all situations where the partition function cannot be evaluated exactly.

### Spot penalty optimization is the most efficient way to recover information

Spot penalty optimization is an iterative process where each iterative step requires the evaluation of some number of series terms. An optimization requiring *t* iterations thus multiplies computation time by a factor of *t* relative to the simple evaluation of the series. Alternatively, one could spend the extra computation time on taking the series to a higher order without spot penalty optimization. Figure 6A plots the unrecovered information when a) taking series 2 to a certain order without optimization, versus b) using spot penalty optimization on only the first term yielding 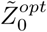 The dotted line in each panel shows the median number of terms requiring the same computation time as 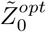 The Toy scenario shows that, if the series expansion is carried deep enough, it becomes more accurate that 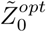: in other words the difference 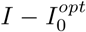 becomes negative. However, for the practical scenarios of Experiments 1 and 2 this crossover point requires taking more terms than would be needed to match the computational cost of calculating 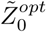 (the dotted line). Based on these results, we recommend always performing spot penalty optimization, especially for larger reconstructions.

**Figure 6:**
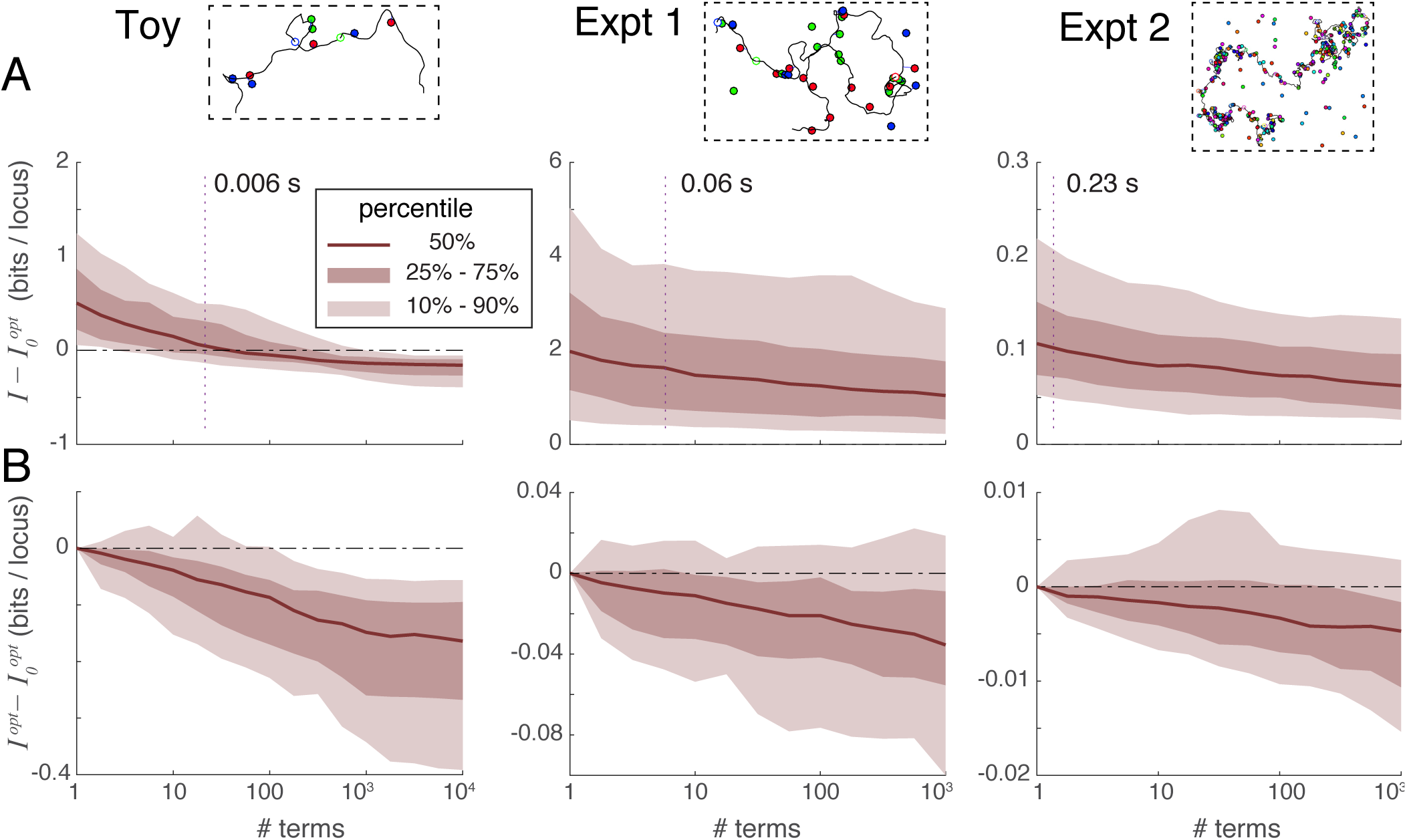
Optimization in conjunction with series expansions. **A.** Comparison of unrecovered information using series expansions without iteration, denoted *I,* to the unrecovered information obtained by optimizing spot penalties using only the first series term, denoted 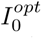, over three experimental situations. Vertical dotted lines indicate the median number of series terms computable with the same computational time as was required to obtain 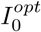. For Experiments 1 and 2 the difference 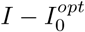 is typically positive at the intersection of the dotted line, indicating that spot penalty optimization method is the more efficient way of recovering information. **B.** Comparison of unrecovered information using spot-penalty optimization in conjunction with multiple series terms versus optimization over 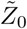, showing the added benefit of including more terms in the series.

### Series expansions can improve optimization information recovery

Although spot penalty optimization is the most efficient way to recover information, that process alone can only extract a certain fraction of the recoverable information: once the cost function is zero, optimization can proceed no further despite the problem not having been solved exactly. At this point, the only way forward is to go higher in the order of series terms used; we can still apply spot penalties to this sum of terms and iteratively optimize them as before using Eqs 1 and 2. Figure 6B plots the difference in unrecovered information when applying spot penalty optimization between a) a variable number of terms in series expansion 2, and b) only 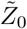 (the first series term). This figure shows that including additional series terms in the optimization improves the information recovery, albeit at a slow rate (especially for large problems).

### 20-color labeling leads to near-perfect reconstructions

As shown in Figure 5A, the unrecovered information for the whole-chromosome Experiment 2 averages around 0.2 bits per locus, implying near perfect mapping probabilities. However, because these results were based on randomly-generated unconfined conformations, they do not establish whether such good information recovery is possible with real chromosomes which are likely to be more compact. To test Experiment 2 on realistic chromosome conformations, we generated four plausible conformations of human chromosome 4 by running the GEM software package [13] on the smoothed human Hi-C data set provided by Ref. [14] and using a 3D spline interpolation to increase the resolution from 1 Mb to 100 kb. These conformations were then virtually labeled at 300 randomly-selected loci with simulated experimental error as before (100/200 nm localization error in xy/z, 10% missing-and extra-spot rates). Mapping proba-bilites were reconstructed by taking series expansion 2 to the lowest order that included at least 1000 terms, then applying and optimizing spot penalties. Compared with the random-walk conformations used to test the Experiment 2 scenario, these reconstructions did somewhat worse (∼ 0.25 versus ∼ 0.2 bits of unrecovered information per locus) owing to fact that physical confinement of chromosomes increases the density of competing spots in the image.

Despite the drop in performance, 0.25 bits of unrecovered information per locus is still an extremely strong reconstruction, implying that the correct locus-to-spot mappings are assigned *p-*values averaging around 2^−0.25^ ≈ 84 % Starting from such accurate and confident mapping probabilities, one can infer a reasonable conformation simply by assigning each locus to the spot to which it maps with the highest probability (or calling a missing spot if 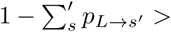 any *P*_*L* →_*_S_*), and drawing a line in the image that connects these spots in genomic order. The conformations produced by this simple rule are shown in Figure 7: the correct conformation is shown with a blue line and errors in the inferred conformation are shown in red. The reconstructed conformations are ∼ 90% accurate as determined by an alignment between the true and inferred spot sequences traveling along the DNA contour (40, 33, 26, 33 mismatches + indels for the 4 respective experiments). Most mistakes are of a sort that does not change the large-scale structure. For example, one common error is to erroneously skip one or more spots in the image (since at present `align3d` overestimates the missing-spot rate), thus ‘looping out’ a small part of the conformation and effectively lowering the resolution

**Figure 7:**
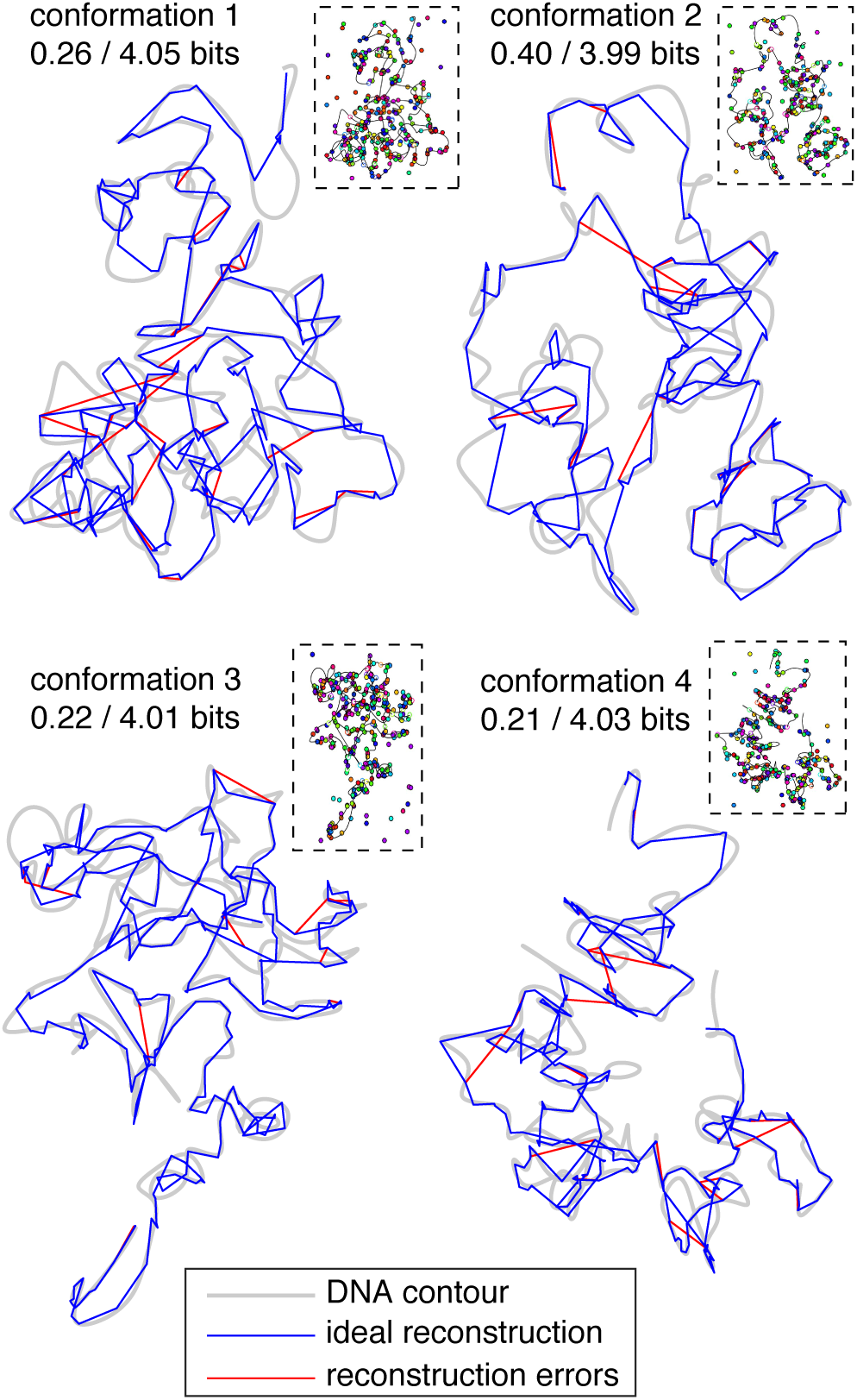
Simulated reconstructions of 4 plausible conformations of human chromosome 4. Grey shaded lines indicate the underlying DNA contours; blue lines trace the ideal reconstructed contours given the simulated labeling (shown in small panels at upper right); red lines show our reconstructed contours where they deviate from the ideal contours. Captions indicate the amount of unrecovered information per locus after/before the reconstruction process.

## Discussion

We have developed and evaluated two improvements to the `align3d` method for reconstructing chromosome structure. Both of these improve the partition function estimates that determine the locus-to-spot mapping probabilities, which can provide the basis for (probabilistic) reconstructed conformations. The first improvement is a more robust spot-penalty optimizer that allows for large-scale reconstructions involving hundreds of labeled loci, such as will be needed to uncover whole-chromosome conformations. The second improvement is two series expansion formulas for the partition functions, which in principle allow the mapping probabilities to be solved to arbitrary accuracy within the limitations of the experiment and the underlying DNA model. In practice, the series approach is difficult for two reasons: 1) there are a huge number of terms in each series expansion, and 2) the lowest-order approximation 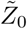 overestimates *Z* by many orders of magnitude, unlike other series expansions where the initial approximation is close to the final answer. Despite the difficulties, the series formulas that we give offer some way forward to improve on the original estimate 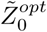. Of the two formulas, we recommend using series expansion 2, which has the larger-magnitude terms and thus recovers the most information when only a few terms can be evaluated.

Our problem of finding likely (i.e. low-free-energy) DNA conformations passing through a set of imaged spots is similar to the well-known traveling salesman problem (TSP), in which a salesman must find the shortest route connecting a set of cities. Somewhat more closely related is a generalization of the TSP called the time-dependent traveling salesman problem (TDTSP) [10], where the intercity distances change every step on the tour; this is analogous to our situation where the free energy needed to thread DNA between two spots depends not only on their separation but also on the length of DNA used to connect them. In our case, the presence of missing and extra spots generalizes our problem still further: in the TDTSP analogy the salesman would be allowed to skip stops and cities for a penalty. Our main finding is that the partition function of this generalized TDTSP (which encompasses traditional TSP and TDTSP problems) can be expressed as a sum of terms computable using a (modified) forward-backward algorithm. This result should also apply to other route-finding applications, and might provide a fresh approach since research has historically focused on estimating the single optimal solution rather than the partition function.

From a genomic standpoint, our most exciting result is that the combination of our computational improvements together with 20-color labeling technology gives almost perfectly-accurate reconstructions. Out of ∼ 4 bits per locus of uncertainty inherent in the reconstruction problem, our method recovers about 3.75 bits, despite somewhat overestimating the missing-spot rate in these simulations. Such confident mapping probabilities allow for the direct construction of individual conformations that are about 90% accurate. We want to emphasize that our reconstructions used model parameters that were correctly calibrated, but not necessarily ‘correct’ in terms of details that an experimenter would not know. For example, because the conformations were generated using a wormlike chain DNA model but `align3d` assumes a Gaussian chain model, the underlying DNA model used in the reconstruction is wrong in a sense, just as it will be using *in vivo* data where the underlying DNA model is unknown. However, we did set the Gaussian decay constant to correctly match the persistence length as that can be measured experimentally. Additionally, the correct average rates of missing and extra spots over all experiments were provided to the analysis, but using those `align3d` still had to estimate the actual number of missing spots per color in each experiment. The robustness of the analysis to experimental unknowns gives evidence that reconstructions using real-world experimental data will be of similar quality to those in our simulations, and if so then direct measurement of chromosome conformations is possible today with current technology.

## Acknowledgments

The authors want to thank Rani Powers and Jenny Mae Samson for helping review the manuscript. This work is supported by the Boettcher Foundation (J.C.C.), NIH grant 2T15LM009451 (B.C.R.), and a Cancer League of Colorado grant.

## Appendices

### Appendix 1: justification for gradient-free weight update rule

The optimization of the spot penalties uses a cost function that is entirely determined by the total probability *P(s)* of mapping a locus to a given spot *s* in the image:

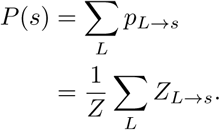

Both *Z*_*L→S*_ and *Z* sum terms that are products of the spot penalties *q*_*s*_, owing to the fact that a direct influence on any one mapping probability indirectly influences all other mapping probabilities. However, when the penalty on some spot *s* is far away from its proper value, any mapping probability *P*_*L*_*→s* to that spot tends to be saturated very close to either 0 or 1. In this case we make the approximation that a given penalty factor *q*_*s*_ only affects any given *P*_*L*_*→s* directly as a multiplying factor, and does not affect the mapping probabilities to other spots, since the indirect influences are small until the mapping probability is out of saturation. Under this approximation *Z*_*L→S*_ ≈ *fs · a*_*s*_, and *NZ =* Σ_*L*_ Σ_*s*_ *Z*_*L→S*_ ≈ *f*_*s*_ · *a*_*s*_ + *b*_*s*_, so

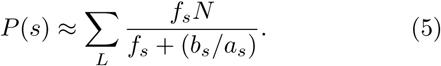

Since our objective is to find an updated 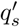 causing the sum of mapping probabilities to be some target *P’ (s*), we also write:

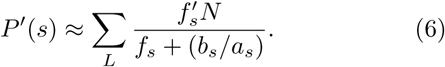

Solving Eqs 5 and 6 together to eliminate the unknown *b*_*s*_*/a*_*s*_ gives us the update rule:

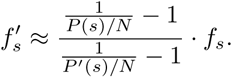

The cost function contains two terms: 1) a penalty for the overall difference between the expected rate of missing spots *p_fn_* versus that inferred from the summed *P(s)*; and 2) a penalty on *P(s)* if it exceeds 1, which would indicate a > 100% likelihood of a locus mapping to that spot. Our current implementation simply makes two updates, one pushing *P(s*) → *p_f_* and the second pushing *P(s) →* 1 for spots violating normalization. Doing so leads to the update rule given by Eq. 1.

### Appendix 2: series expansion derivations

Here we prove a) each series converges to the true partition function when all terms are counted, and that b) a given truncation of each series using our recommended selection of series terms counts every possible legal or overlapping conformation zero or more times (despite many series terms having negative coefficients), and therefore produces positive mapping probabilities. Notice that both expansions count all legal (non-overlap ping) conformations once from 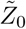, and the goal of considering higherorder series terms is thus to minimize the weight of the illegal overlapping conformations without their weights ever becoming negative. We note that conformations overlapping at adjacent loci are automatically eliminated from all terms in the calculation, so the individual conformations we consider here are assumed to overlap themselves only between non-adjacent loci.

For our proofs we will define an illegal overlap as a set of loci in an illegal conformation mapping to a given spot in the image. Illegal overlaps are to illegal conformations as illegal constraints are to higher-order series terms, and we use the same set notation for overlaps as we do for constraints.

Our derivations make repeated use of the following identity, taken from Eq. 3.1.7 in Ref. [15].

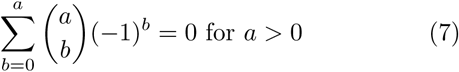

This relation follows from the fact that the left-hand side is a series expansion for (1 – 1)^*a*^.

#### Series expansion 1

A given conformation bearing a set of illegal overlaps *θ* is counted by each constrained partition function whose set of illegal constraints (indices) are a subset of *θ.* Thus the weight *W* _*θ*_ of this conformation in the full partition function *Z* is:

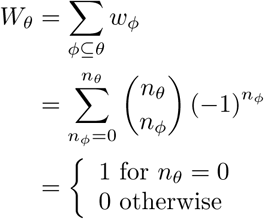

where we have used Eq. 7 in the final step. This result shows that series expansion 1 counts only conformations having no overlaps, i.e. for which *n* _*θ*_ = 0.

Suppose that one follows our prescription for evaluating a subset of terms, namely all terms that are a subset of overlaps *ψ*. Then using the same formula, we reason that a conformation with overlaps *θ* ill be given the following weight:

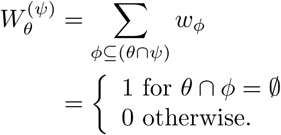

Thus our selection of series terms for expansion 1 eliminates all conformations from the partition function having any overlaps contained in our set *ψ.*

#### Series expansion 2

Expansion 2 allows unconstrained loci to revisit spots that already have constrained loci. In this case, a conformation whose set of nonadjacent overlaps is denoted *θ* will be counted by a partition function having overlaps *ϕ* if each individual overlap *ϕ*_*i*_ is *contained in* an individual overlap *θ*_*i*_, in the sense that *φ* _*i*_ maps any subset of 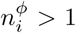 loci in *θ* _*i*_ to the same spot as *θ*_*i*_. Then the weight of this conformation in the expansion is

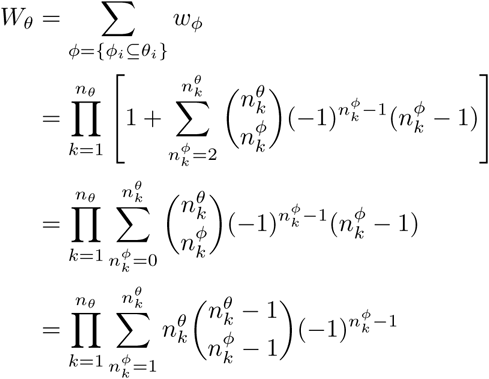

where the fact that 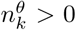 allowed Eq. 7 to eliminate a term in the last line. Defining 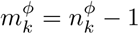 gives us:

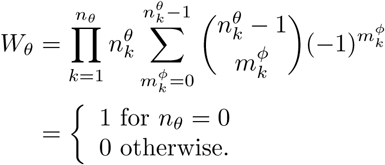

again using Eq. 7. Therefore the complete series counts only non-overlapping conformations.

Our prescription for choosing series terms is to select a set of single-locus-to-spot mappings Ψ,and generate all series terms whose illegal overlaps are built only from mappings in Ψ To be formal, we will compile all locus-to-spot-*s*-mappings in Ψ for each given spot *s* (assuming there are 2 or more) into a single overlap, and use ψ to denote the set of these overlaps: then the series order *Nψ* is the sum of the number of loci over all elements *ψi.* Our rule is to include all series terms *ϕ* whose illegal overlaps are built entirely from *subsets of ψ,* in the sense that a given overlap to spot *s* contains a subset of the loci in the element *ψ i* that maps to spot s. For example if ψ contains the element (*ACE*)_*s*_ then ϕ ⊇ {(.*AC*)_s_, (*CE*)_*s*_, (*AE*)_*s*_, (*ACE*)_s_}. Using these rules, a given conformation having illegal overlaps *θ* will be given the following weight in the expansion:

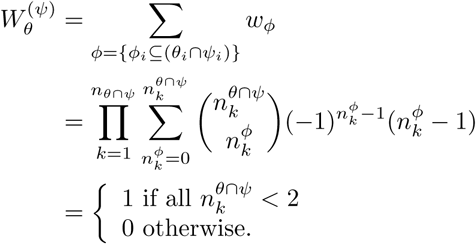

Thus series 2 eliminates all conformations having any overlap (*L*_1_,*L*_*2*_,…) → *s* where *L*_1_ *→ s* and *L*_*2*_ *→ s* are contained in Ψ.

### Appendix 3: Supplementary figures

**Figure S1:**
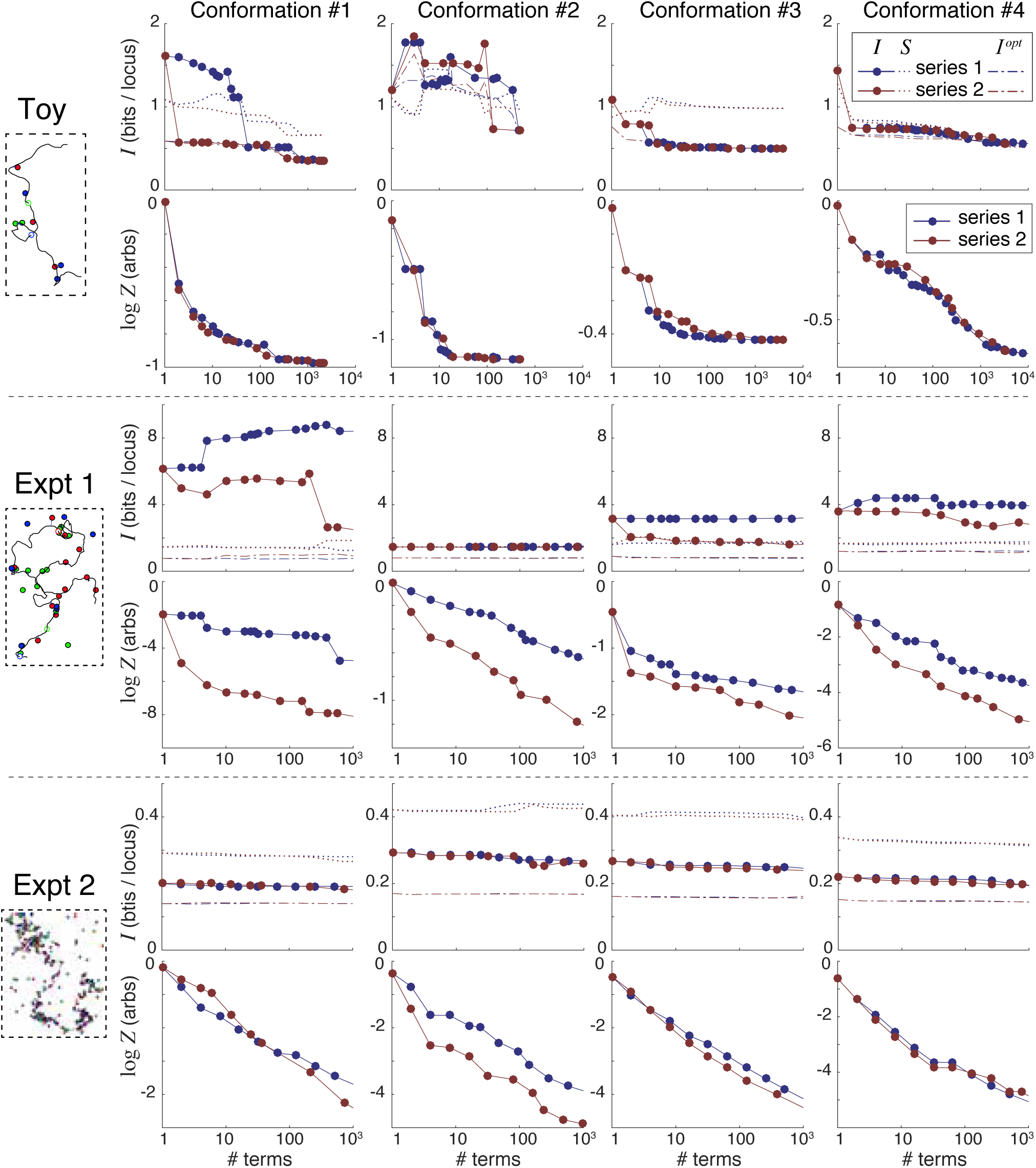
Information recovery from individual simulations. Conformation #1 of each series is shown at far left. Panels to the right show unrecovered information (*I*), entropy (*S*) and log *Z* as a function of the number of series terms included. Dot-dashed lines show the unrecovered information of 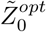 using optimized spot penalties on top of the given number of the series terms.

**Figure S2:**
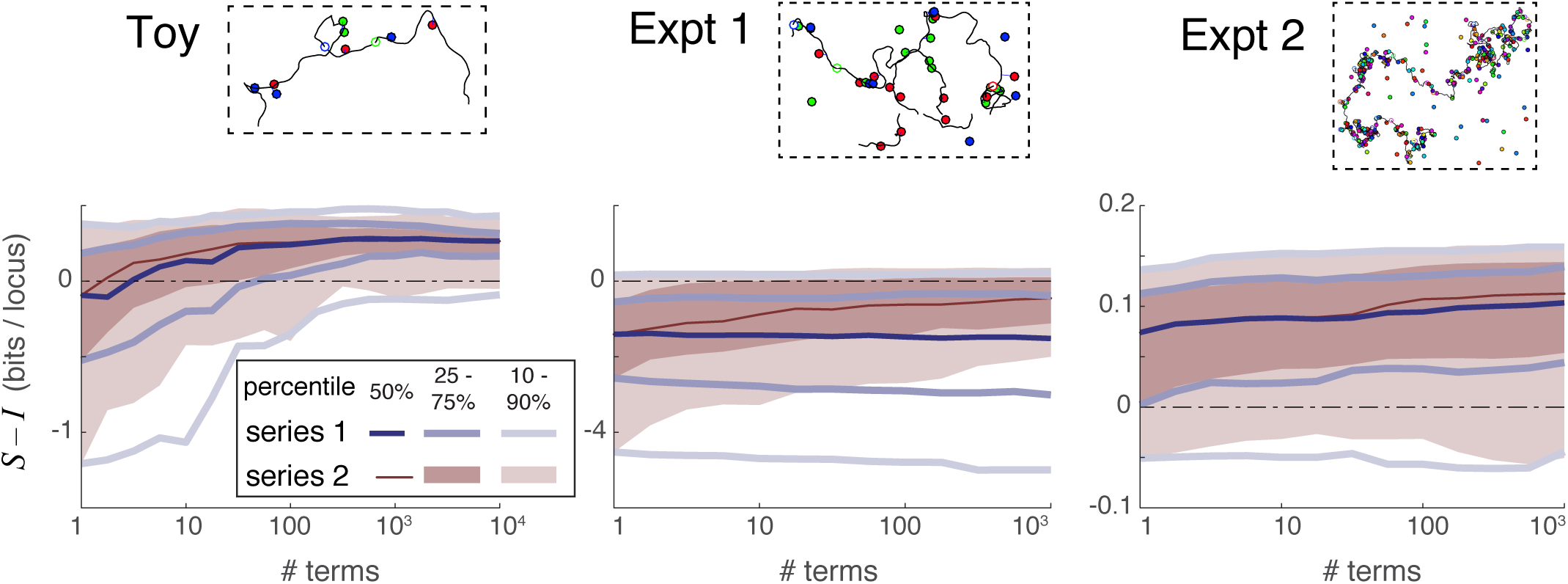
Accuracy of entropy as a proxy for unrecovered information. Distributions showing the difference between entropy *S,* which is a blind estimate of unrecovered information, and actual unrecovered information *I* in each of the 3 simulation scenarios considered, as a function of number of series terms. No spot penalties were used for these results. Each distribution shown encompasses the *S − I* curves of all 100 simulated reconstructions in one scenario.

**Figure S3:**
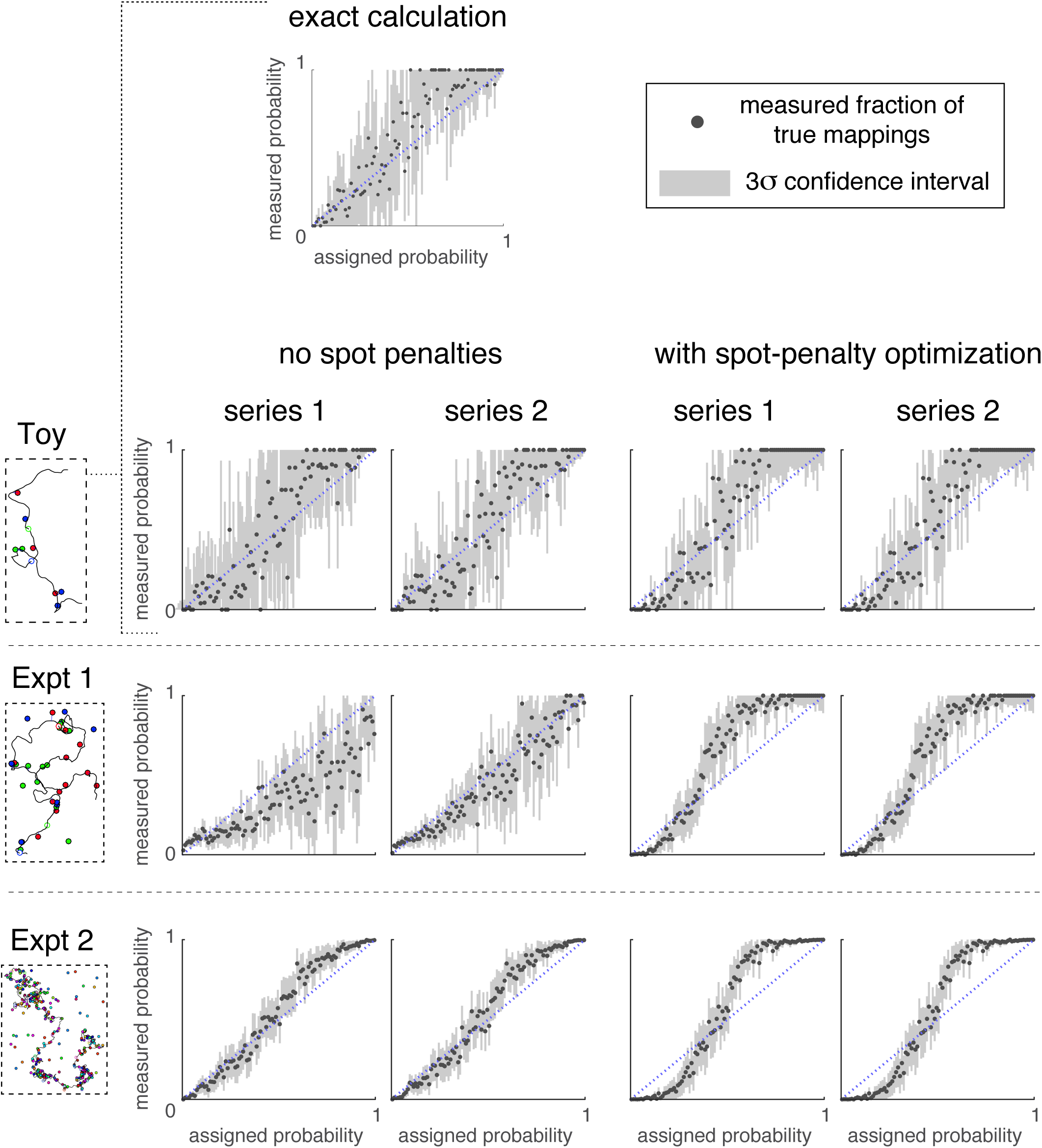
Accuracy of mapping probabilities. Binned mapping probabilities (x axis) versus the fraction of true mapping probabilities in each bin (y axis), averaged over the various simulated experiments in each experimental scenario. Grey shaded regions show the 3σ range of uncertainty due to counting error.

**Figure S4:**
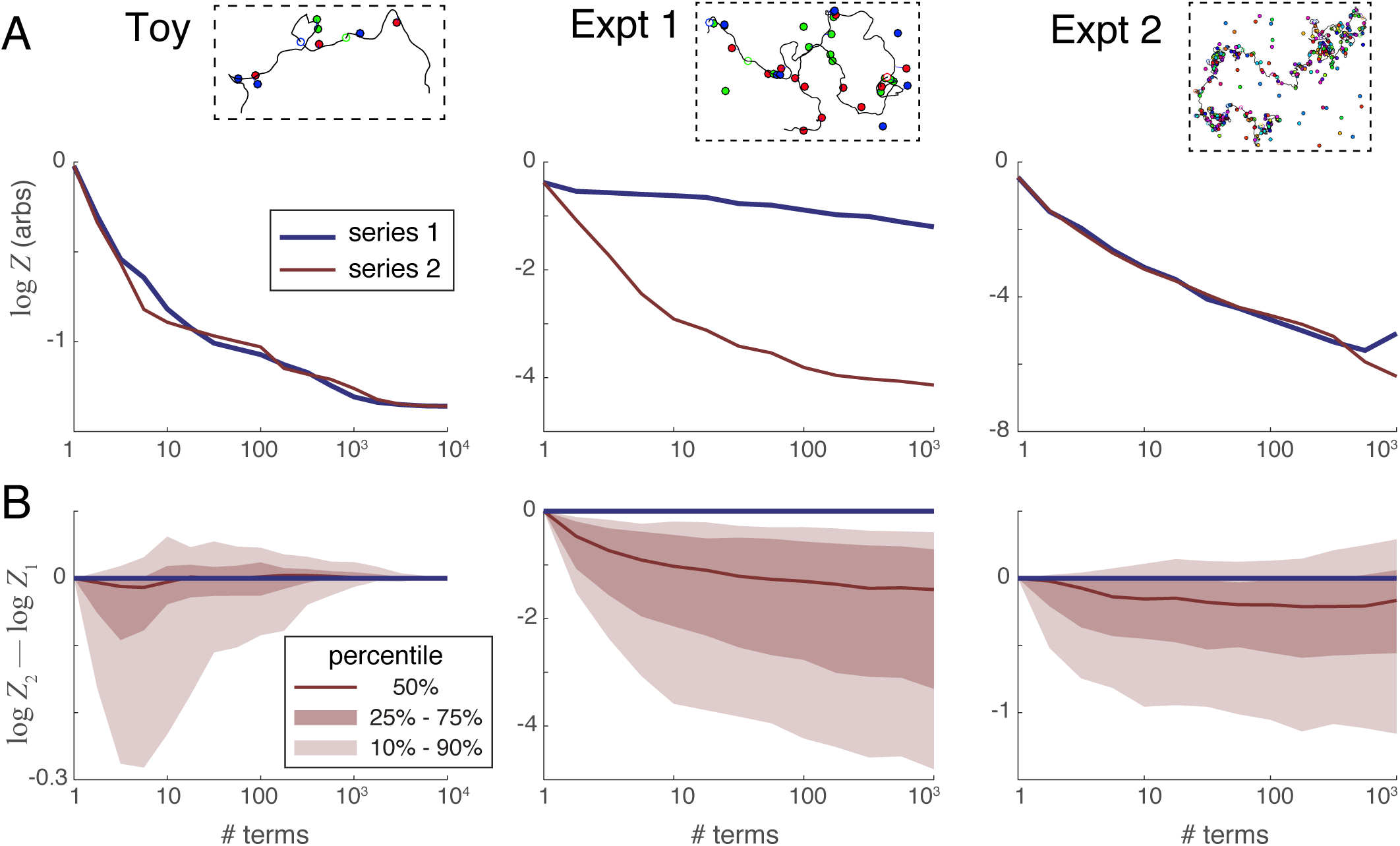
Partition function *Z* versus number of series terms. **A.** Relationship between log *Z* and the number of series terms for each simulated scenario, calculated as a median average of the relationships found in each of the 100 individual simulations in each scenario. **B.** Distributions of the difference between log *Z* calculated using series 2 and series 1. The fact that this quantity is generally negative when some but not all series terms are included shows that series 2 recovers information (i.e. removes unrecovered information *I*) faster than series 1.

## Footnotes

1 Depending on how the experiment is done, two spots of the same color sufficiently close in the image may appear as a single spot where the conformation self-overlaps. We prefer to treat this scenario as a missing-spot measurement error rather than relax the one-spot-per-locus rule. If the spots have been properly localized, then the underlying conformation visits any given spot once at most.

